# Context-dependent decision-making in the primate hippocampal-prefrontal circuit

**DOI:** 10.1101/2023.09.08.556931

**Authors:** Thomas W. Elston, Joni D. Wallis

## Abstract

What is good in one scenario might be bad in another. Despite the ubiquity of such contextual reasoning in everyday choice, how the brain flexibly utilizes different valuation schemes across contexts remains unknown. We addressed this question by monitoring neural activity from the hippocampus (HPC) and orbitofrontal cortex (OFC) of two monkeys performing a state-dependent choice task. We found that HPC neurons encoded state information as it became available and then, at the time of choice, relayed this information to OFC via theta synchronization. During choice, OFC represented value in a state-dependent manner: many OFC neurons uniquely coded for value in only one state but not the other. This suggests a functional dissociation whereby HPC encodes contextual information that is broadcast to OFC via theta synchronization to select a state-appropriate value subcircuit, thus allowing for contextual reasoning in value-based choice.

## Introduction

Cognitive maps provide us with an internal representation of relationships and structure in our environment. They serve as mental blueprints, enabling us to remember locations, plan efficient routes, and generate novel plans on the fly. Although cognitive maps are famously associated with hippocampal place cells and spatial navigation (*1*), recent advances point to a more generalized cognitive map theory that extends beyond spatial cognition to incorporate non-spatial domains such as value, conceptual knowledge, and abstract thought (*2-5*). Recent studies suggest that the orbitofrontal cortex (OFC) also encodes a cognitive map (*6-10*). However, single neuron recordings in monkeys show that OFC typically encodes little information about sensorimotor contingencies (*11-15*) and it may be that OFC is primarily involved in utilizing rather than constructing cognitive maps. This contrasts with HPC where neurons encode sensorimotor contingencies, in addition to spatial and temporal contexts, precisely the kind of information that is essential for building an internal model of the task at hand (*16-18*). HPC densely innervates OFC (*19, 20*) and interactions between them via theta waves have recently been implicated in value-based choice (*21*). Here, we tested the hypothesis that both HPC and OFC make critical contributions to map-driven behavior, with HPC responsible for constructing a cognitive map which provides information about task state to the OFC to generate reward predictions that can be used to guide flexible decision-making.

## Results

To test this hypothesis, we trained two monkeys (subjects K and D) to perform a state-dependent decision-making task. The subjects were required to choose between reward-predictive pictures where the amount of reward associated with the picture depended upon a task state (Fig. 1A). On each trial, subjects were cued as to whether the current trial should be evaluated according to the value scheme of state A or B. Then, after a brief delay, they were presented with either one (forced choice, 20% of trials) or two options (free choice, 80% of trials) to select from. Critically, the picture values on each trial depended on the cued task-state. Therefore, to perform the task, the subjects flexibly updated the values assigned to the choice options and used these flexible, state-dependent valuations to guide their responses. Both subjects were proficient at the task and were equally accurate across both task states (subject K: state A correct trials = 98 ± 0.3 %, state B = 98 ± 0.4 %, paired *t*-test, *t*(4) = 1.8, *p* > 0.1; subject D: state A = 95 ± 0.9 %, state B = 93 ± 1.0 %, *t*(9) = 1.7, *p* > 0.1).

**Figure 1.**
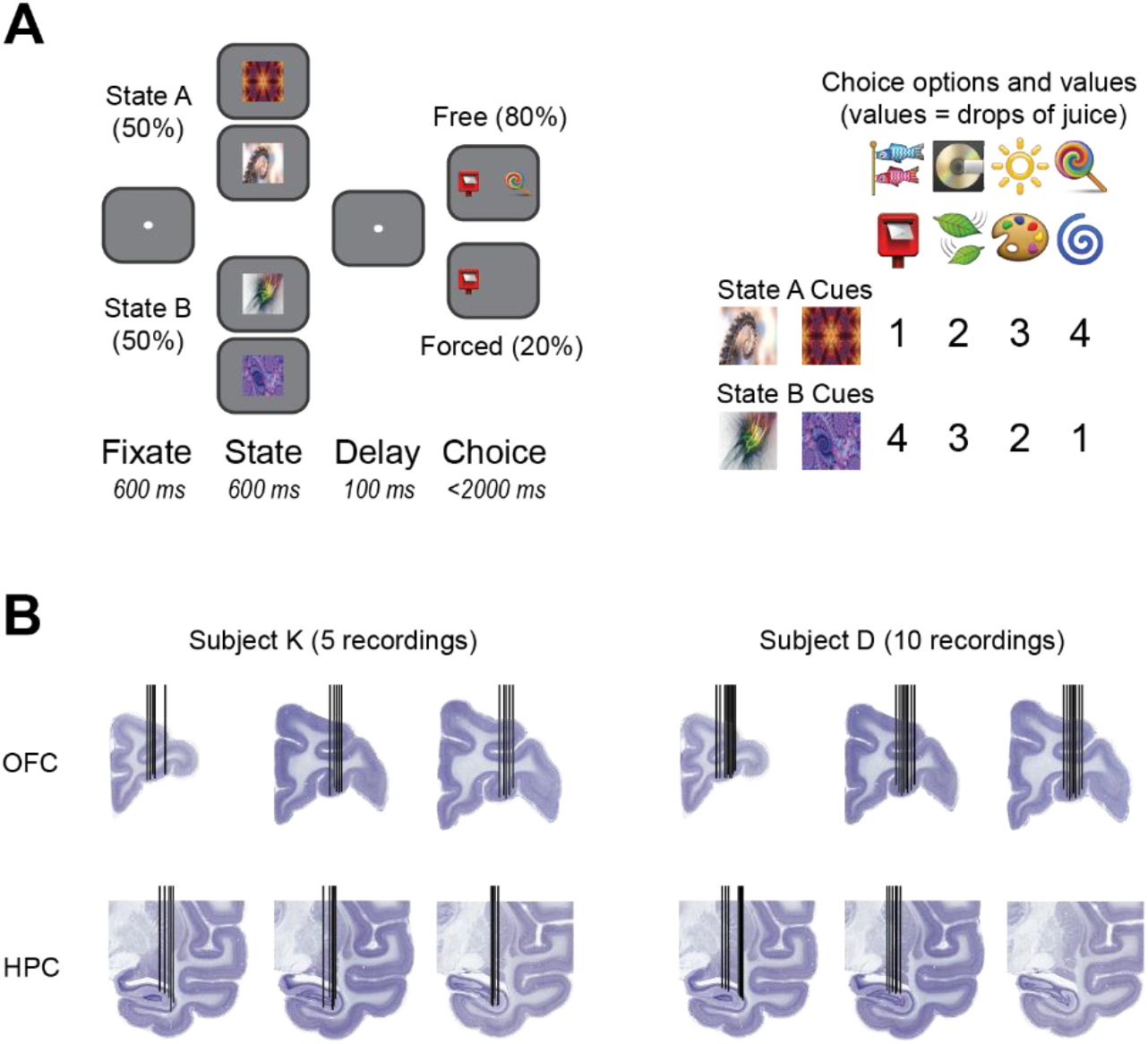
Behavioral task and recording locations. A) Structure of the state-dependent choice task. Subjects initially fixated and were then shown one of four state cues (two per state). After a delay, subjects were presented with either one (forced choice) or two (free choice) choice options. The optimal choice depended on the state cued earlier in the trial. States were varied pseudo-randomly across trials. To unconfound neuronal activity related to the physical properties of the stimuli from the meaning they signified, we used two distinct cues for each state and value. B) Electrode trajectories and recording locations for each subject overlayed on representative Nissl-stained sections from the NIH Blueprint NHP Brain Atlas (blueprintnhpatlas.org). Each black vertical line indicates a single electrode trajectory.

To understand the neural mechanisms of state-dependent valuation and choice, we simultaneously recorded neurons in HPC (K = 179 neurons, D = 125 neurons) and OFC (K = 251 neurons, D = 281 neurons) as subjects performed the task (Fig. 1B). We assessed the tuning of individual HPC and OFC neurons during different task phases using linear regression models (see Methods). Across both subjects, we found that HPC neurons were more likely to encode state information than OFC neurons during the state epoch whereas the reverse was true during the choice epoch (Fig. 2). These results reflect a functional double-dissociation of state-encoding, suggesting that HPC initially encodes state when state information becomes available whereas OFC encodes state information when it must be used to guide a decision.

**Figure 2.**
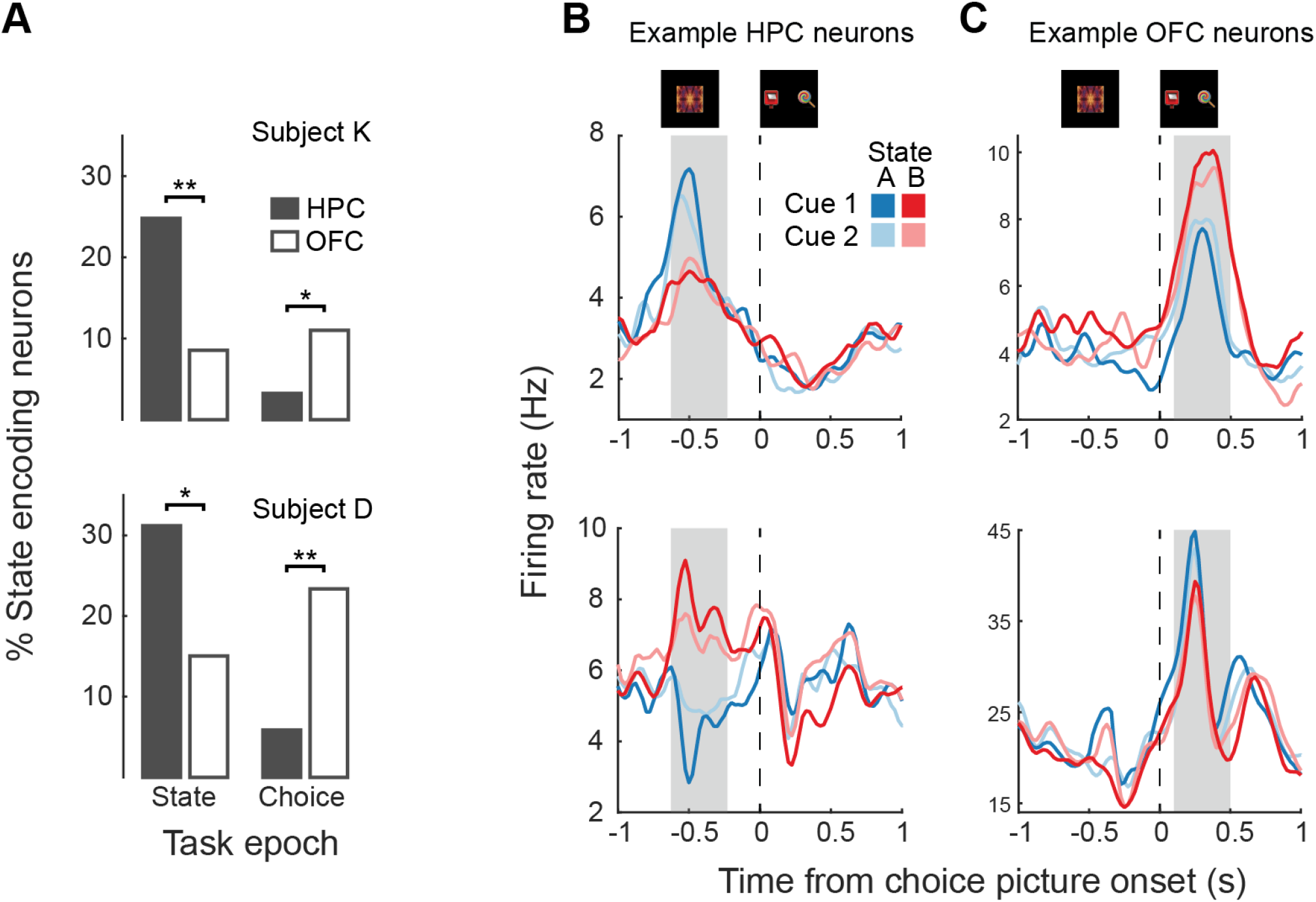
Neuronal encoding of state in HPC and OFC. A) Prevalence of state-encoding neurons in HPC and OFC during the state and choice epochs. * = *p* < 0.05, ** = *p* < 0.005, *χ*^2^ test. B) Example HPC neurons that significantly encoded state information during the state epoch. The top neuron responded more strongly to State A, while the bottom neuron responded more strongly to state B. The shaded region indicates the fixed window during the state epoch firing rates were analyzed in. C) Example OFC neurons that significantly encoded state information during the choice epoch. The top neuron responded more strongly to State B, while the bottom neuron responded more strongly to state A. The shaded region indicates the choice epoch.

Our single neuron results suggest that HPC initially encodes state-information and then relays it to OFC so that state can inform cortical valuation and decision representations. HPC strongly projects to OFC (*19*) and HPC-OFC interactions in the theta band (4-8 Hz) have previously been implicated in updating OFC value representations (*21*). Given that neuronal state-encoding was strongest in HPC during the cue phase but then shifted to OFC during the choice phase, we wondered whether the structures could be communicating. Across both subjects and brain areas, theta responses were visible on nearly every individual trial, and they appeared to align to specific task events (Fig. 3A). We quantified this effect via a cross-trial phase-alignment measurement that measured the mean resultant vector length, *R*, of phase angles across trials. In both subjects and both brain areas, theta phases showed strong cross-alignment at the time of fixation and at the presentation of the state cue (Fig. 3B). Interestingly, we did not observe a distinct response to the onset of the choice options, showing that these responses are not simply sensory-driven events.

**Figure 3.**
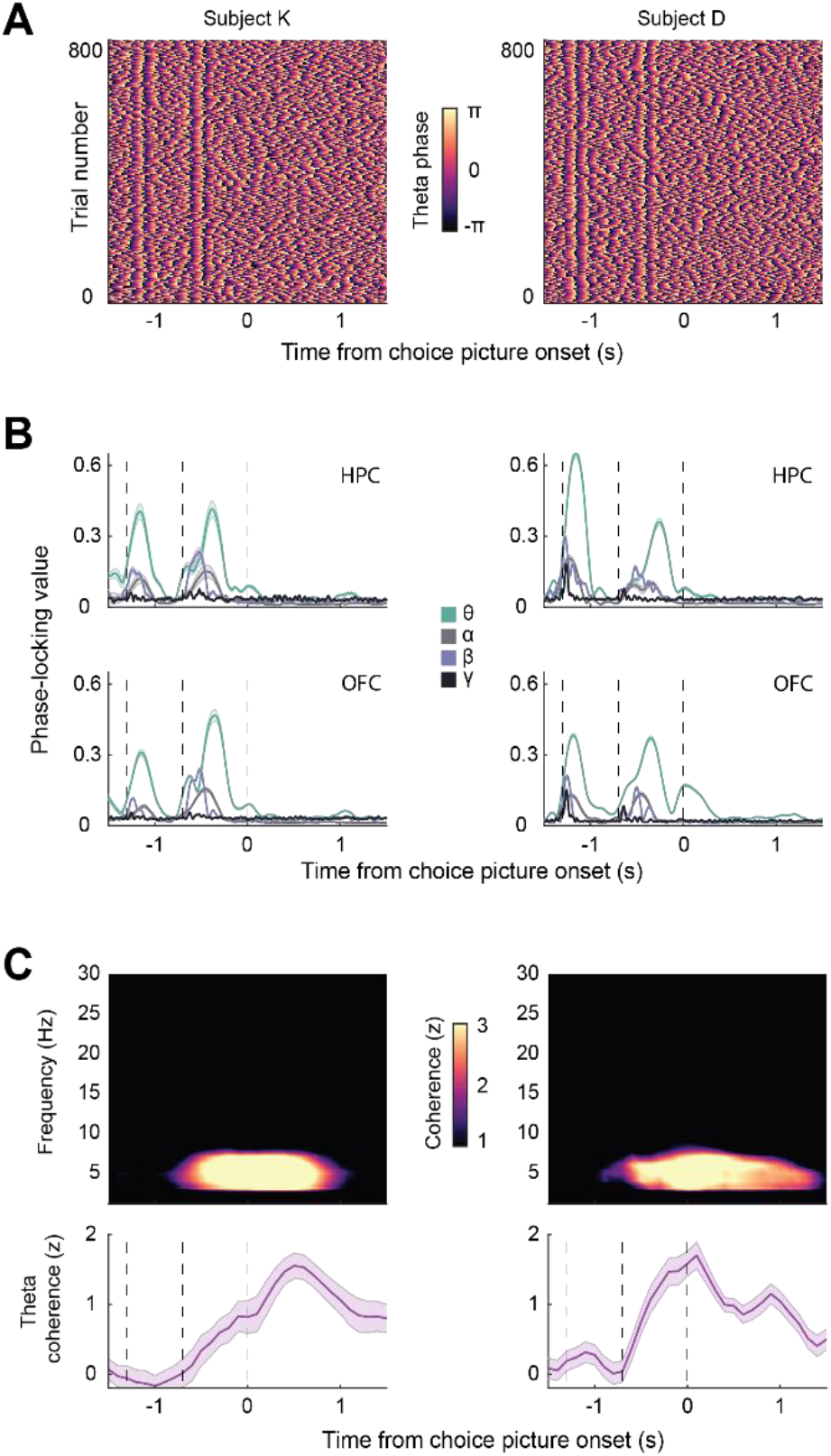
Theta activity in HPC and OFC. A) HPC theta phase over the course of the trial from a representative recording session from each subject. Each row represents the instantaneous phase at each of 800 trials within a single session on a single HPC electrode. B) Cross-trial phase alignment at theta (4–8 Hz), alpha (9-12 Hz), beta (13–30 Hz), and gamma (30–60 Hz) frequencies. Error bars represent bootstrapped 95% confidence intervals. The dashed vertical lines indicate, from left to right, the onset of the fixation cue, state cue, and choice cue. Theta exhibited the strong phase alignment at fixation and the presentation of the state cue, but not the presentation of the choice options. C) HPC-OFC coherence in each subject. The top row shows session-mean coherences for HPC-OFC electrode pairs in each subject. The bottom row shows the time course of the mean theta coherence, which increased shortly after the onset of the state cue and peaked at the time of choice. Error bars denote bootstrapped 95% confidence intervals. The dashed vertical lines indicate, from left to right, the onset of the fixation cue, state cue, and choice cue.

Having established the presence of theta oscillations within our task, we next measured HPC-OFC communication via analysis of oscillatory coherence (*22*). Theta coherence increased shortly after the onset of the state cue and peaked after the onset of choice options (Fig. 3C). To quantify the direction of information flow between OFC and HPC, we used generalized partial directed coherence (GPDC; see Methods). Across both subjects, we measured stronger bottom up, HPC→OFC directionality than OFC→HPC directionality (subject K: net directionality = 0.1 ± 0.007 95% confidence interval; subject D: 0.07 ± 0.01, *p* < 0.001 in both subjects, bootstrapping procedure).

We next examined whether HPC and OFC neurons were significantly modulated by LFP oscillatory activity during the state-cue period. Many neurons in both brain areas and subjects were significantly theta-modulated (subject K: OFC = 81/351 or 23%, HPC = 40/179 or 22%; subject D: OFC = 136/231 or 59%, HPC: 64/125 or 51%) whereas the proportion of neurons modulated by other frequency bands did not exceed chance levels (binomial test, *p* > 0.1 in all cases). We found that most theta-modulated neurons in HPC tended to fire during the rising phase of the theta oscillation (Fig. 4A, B), consistent with prior findings in humans (*23*). Thus, the ongoing theta rhythm appears to organize the firing of HPC units during the period when theta coherence occurs between regions. In addition, the first theta peak during the state epoch occurred significantly earlier in HPC than OFC in both animals (Fig. 4C) supporting the GPDC analysis.

**Figure 4.**
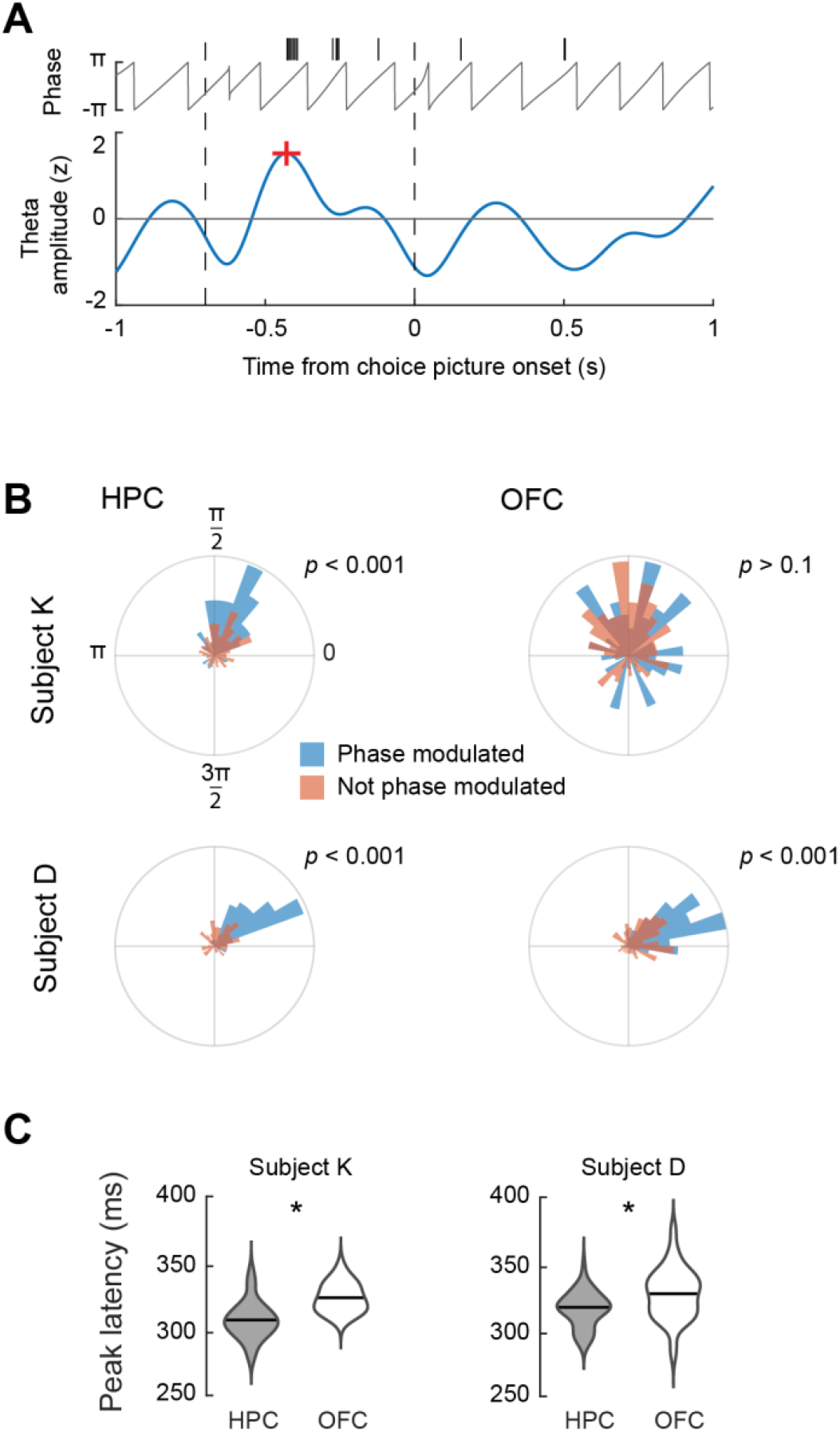
Phase-modulation of single neurons. A) Example of phase locking from a single neuron and a single trial. Black tick marks indicate spikes. The red + denotes the first theta peak. This example is from HPC in subject D. B) Mean angles of neurons that were (blue) and were not (orange) significantly phase modulated by the theta rhythm. HPC neurons in both subjects tended to fire during the rising phase of theta. Results were inconsistent across animals in OFC. The *p* values are from Rayleigh’s test of circular uniformity on the mean angles of the significantly phase-modulated units. C) Mean time of the first peak of the theta rhythm following the onset of the state cue. The first theta peak occurred significantly earlier in HPC relative to OFC in both subjects. * = *p* < 0.001, two-sample *t*-test.

Our final analysis examined how value coding was influenced by task state. For each neuron during the choice epoch, we regressed its firing rate against the value of the chosen option. We fit two such regression models per neuron (one for each state). In both areas, we observed many neurons that encoded value in at least one state (subject K: 48/179 or 27% HPC, 126/351 or 36% OFC; subject D: 54/125 or 43% HPC, 126/281 or 45% OFC). To understand whether the value coding varied by state, we correlated the resulting beta weights. If value coding was independent of state, we would expect a significant, positive correlation of the value beta weights from the two regressions. Indeed, this was the case in HPC but not OFC (Fig. 5A). Furthermore, the correlation was significantly stronger in HPC than OFC (*p* < 0.001 in both subjects; Fisher’s *r* to *z* transform and test). Thus, OFC encodes state-dependent values, whereas the value coding in HPC is independent of state.

**Figure 5.**
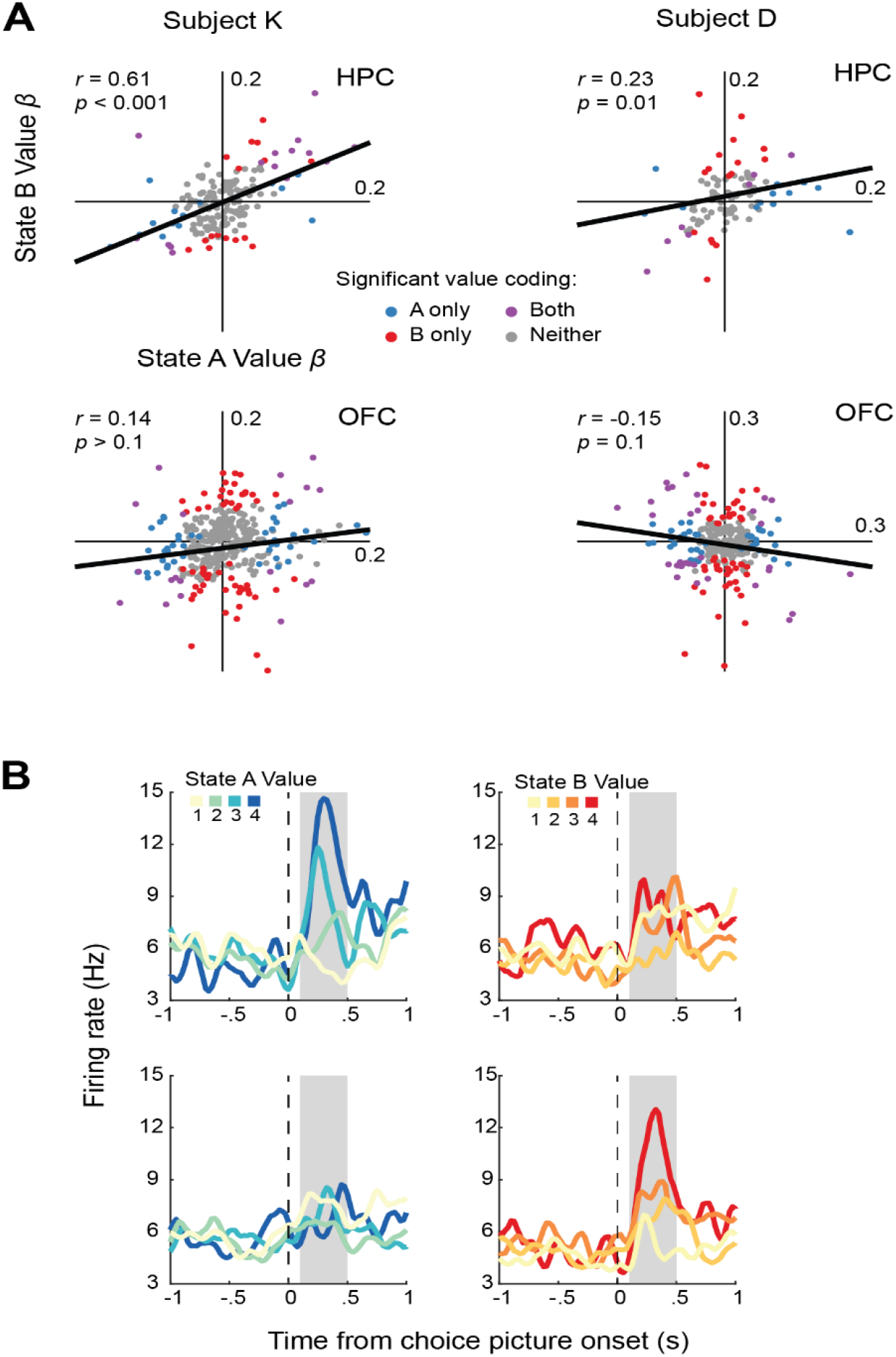
OFC encodes state-dependent values. A) Value beta weights for each neuron in each state across subjects and brain areas. Each data point is a single neuron. The *r* values describe Pearson correlations. B) Example OFC neurons that encoded value in only state A (top) or B (bottom). The shaded region indicates the choice epoch.

## Discussion

Our results suggest that both HPC and OFC are central to behaviors that depend on cognitive maps: HPC neurons group cues into behaviorally relevant categories that then select state-appropriate decision circuits in the OFC via theta synchronization. More generally, our results support a division of labor between HPC and OFC (*24*), whereby HPC encodes conceptual task states (*18, 25*) and OFC uses those states to inform valuation and choice. One interpretation as to why OFC formed distinct state-subcircuits could be to facilitate credit assignment during learning. Indeed, recent results suggest that OFC is critical for learning to attribute specific outcomes and expectations to specific task states (*9, 26*). Forming distinct value-representations across states could also mitigate representational conflict when generating reward predictions. For instance, in our task, the stimuli associated with low value in state A are also associated with high value in state B. By creating distinct subcircuits, the OFC may be able to reduce the conflict between these opposing value predictions during deliberation. In this way, state-specific value subcircuits could facilitate both initial learning and subsequent cognitive control.

It is increasingly evident that the HPC plays a prominent role in reward-guided behavior. Studies of single neuron activity have demonstrated that HPC may play an important role in goal-directed behavior and reward expectation. In rodents, investigators have decoded information from place cells to show that HPC neuronal activity tends to be biased towards goal locations (*27, 28*). In monkeys, HPC neurons have been found to encode specific value relationships between choice options (*3*). Causal manipulations of HPC also impair value-based decision-making, as shown by damage to HPC in humans (*29*) or electrical microstimulation of HPC in monkeys (*21*).

In contrast to HPC, OFC has a long history of being associated with reward processing (*11, 30-32*). However, these studies have frequently emphasized that OFC abstracts values across choice options to derive a common neural currency for value (*11, 13, 33, 34*). In contrast, our results suggest that the OFC value code is much more flexible than previously described and can be flexibly reconfigured on the fly. The calculation of these contextually appropriate values requires the coordinated interaction between prefrontal and hippocampal circuits. A range of neuropsychiatric disorders are characterized by dysfunction of these circuits (*35, 36*). A better understanding of the function of these circuits and the principles that coordinate information flow between them could improve treatments that attempt to normalize activity within the circuit (*37*).

## Acknowledgements

This work was funded by NIMH R01-MH117763 and NIMH R01-MH121448 to JDW. The funders had no role in study design, data collection and analysis, decision to publish or preparation of the manuscript.

## Author contributions

TWE designed the experiment, and collected and analyzed the data. TWE and JDW wrote the manuscript. JDW supervised the project.

## Competing financial interests

The authors declare no competing interests.

## Methods

### Experimental model and subject details

All procedures were carried out as specified in the National Research Council guidelines and approved by the Animal Care and Use Committee at the University of California, Berkeley. Two male rhesus macaques (subjects D and K, respectively) aged 8 and 4 years, and weighing 11 and 9 kg at the time of recording were used in the current study. Subjects sat head-fixed in a primate chair (Crist Instrument, Hagerstown, MD). Eye movements were tracked with an infrared system (SR Research, Ottawa, Ontario, CN). Stimulus presentation and behavioral conditions were controlled using the MonkeyLogic toolbox (Hwang et al., 2019). Subjects had unilateral recording chambers implanted that allowed access to both HPC and OFC.

### Task design

Stimuli were presented on a computer monitor positioned at a viewing distance of approximately 30 cm. The subjects were trained to perform a state-dependent decision task. This required them to choose between eight reward-predictive pictures, where the reward amounts associated with the pictures depended on a cued task state (**Fig. 1A**). Subjects self-initiated trials by fixating on a white dot for 600 ms. After initial fixation, the subjects were shown a state cue that indicated whether the upcoming choice should be evaluated according to the value scheme associated with state A or B. After 600 ms, the state cue disappeared, and the subjects had to maintain central fixation for a further 100 ms. Finally, the choice options were presented. On 80% of the trials two options were shown (free choice trials), while on the remaining 20% of the trials there was only one option (forced choice trials). The inclusion of forced choice trials ensured that subjects experienced all reward contingencies. Subjects reported their response by shifting their gaze to their selection and fixating for 300 ms. The values of the choice options corresponded to different amounts of apple juice reward.

To illustrate how state modulates value, and thus choice, consider a trial where the state cue was an orange fractal (indicating state A), and the choice options were the lollipop and postbox emojis. In state A, selecting the lollipop yielded four drops of juice whereas selecting the postbox yielded only one drop of juice. However, had a state B cue been shown earlier (e.g., a purple fractal), selecting the postbox would have yielded four drops of juice and the lollipop only one drop. State cues were pseudorandomized across trials, meaning that there was no serial dependence between trials. This design precluded the use of trial-and-error learning and the optimal decision entirely depended on a given trial’s cued state. Subjects never choose between options with the same value; all other combinations of stimuli were used.

To unconfound neuronal activity related to the physical properties of the stimuli from the meaning they signified, we used two distinct cues for each state and value. Thus, we used two cues to indicate each state (four total state cues) and two pictures to indicate each possible value level (eight total choice options).

### Neurophysiological recordings

Subjects were fitted with titanium head positions and imaged in a 3T scanner. The resulting images were used to generate 3D reconstructions of each subject’s skull and brain areas of interest. We then implanted custom, radiotranslucent recording chambers made of polyether ether ketone (PEEK; Gray Matter Research, Bozeman, MT). Chambers were placed to allow neurophysiological recordings from OFC and HPC.

Each recording session, we acutely lowered up to six Plexon V-(Plexon Inc., Dallas, TX) distributed between OFC and HPC. Each probe had either 32 or 64 contacts per probe with either 50 or 100 μm intercontact spacing. We used OnShape (PTC, Cambridge, MA), a browser-based computer-assisted drawing software to define electrode trajectories and to design custom microdrives to acutely lower the electrodes. The custom drive assemblies were 3D printed using Rigid4000 Resin and a Form3+ printer (FormLabs, Cambridge, MA).

Neuronal signals were acquired and digitized using a Plexon OmniPlex acquisition system with a digital head stage processor. Neuronal signals were acquired at 40 kHz and LFPs were acquired at 1000 Hz. Neurons were identified via manual spike sorting (Offline Sorter, Plexon Inc., Dallas, TX) and retained if their mean firing rate over the recording session was at least 1 Hz.

### Single-neuron tuning

For each neuron, we calculated its mean firing rate, *F*, on each correct trial during two task epochs: the state epoch, defined as the 400 ms period that began 100 ms after the presentation of the state cue, and the choice epoch, defined as the 400 ms period that began 100 ms after the presentation of the choice options. To identify neurons that encoded the task state, for each neuron and each task epoch we performed the following linear regression:

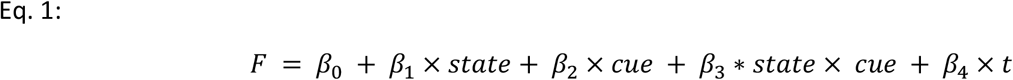

where *state* was a dummy coded variable that indicated which state was cued (A or B), *cue* was a dummy coded variable that indicated the visual properties of the cue (object or fractal), and *t* was a nuisance parameter that indicated the trial number and was used to account for non-stationarity in the neuronal recordings. Within each task phase, neurons were considered state-encoding if *β*_1_ was significant assessed at *p* < 0.05.

To identify how neurons encoded value information, for each neuron we performed the following linear regression on its activity during the choice epoch:

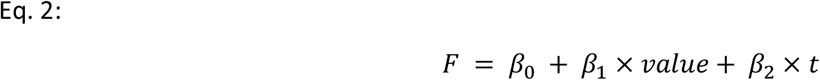

where *value* indicated the number of drops of juice associated with the chosen option, and *t* was a nuisance parameter that indicated the trial number and was used to account for non-stationarity in the neuronal recordings. We performed this analysis separately for trials in which state A was in effect versus state B.

### LFP phase alignment and interareal coherence

To measure cross-trial phase alignment, we extracted phase information from bandpass-filtered LFP signals using the Hilbert transform. Phase-alignment strength was determined by calculating the mean resultant vector length, *R*, across trials:

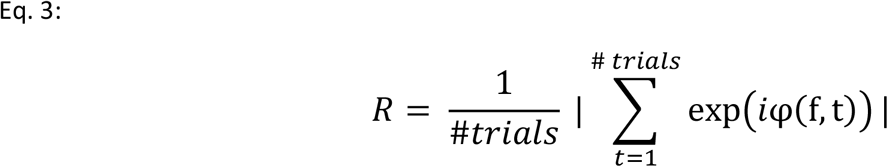

For each recording channel, *i*, this measures the degree to which a given phase, *φ*, of LFP frequency, *f*, is aligned across trials, for each time point, *t*, in the trial. This analysis was done separately for each recording site that had at least one well-isolated neuron (K: HPC = 179 channels, OFC = 351 channels, D: HPC = 125 channels, OFC = 281 channels).

Time-resolved LFP power and inter-regional coherence were measured via multi-taper coherograms (*38*) which used 3 tapers, a 1 s time window, and 90% overlap between windows. In multi-taper spectral analysis, the ‘tapers’ are a set of orthogonal or nearly orthogonal functions (also known as ‘Slepian functions’) that are applied to the time series data before estimating the power spectral density. The Hamming window that is often used in the Fast Fourier Transform (FFT) is an example of a taper. These tapers are designed to reduce spectral leakage and enhance the precision of time-resolved spectral estimates. By applying multiple tapers with slightly different spectral characteristics, the multi-taper approach to signal processing allows for a more accurate estimation of the power spectral density and, in the case of coherograms, coherence between two signals.

We observed a high degree of redundancy of the LFP signal across the contacts of a single V-probe. To prevent oversampling the same underlying signal, we used a single channel per V-probe. For each probe, we identified the channel that had the highest cross-trial phase alignment and at least one neuron. We then computed HPC-OFC coherences between this limited subset of channels. This restricted our analysis to 42 HPC-OFC channel pairs for subject K and 28 HPC-OFC channel pairs for subject D. To facilitate comparison across sessions, coherences were z-scored for each HPC-OFC electrode pair.

To assess the direction of HPC-OFC communication we calculated the GPDC, a frequency-resolved estimate of Granger causality which uses multivariate autoregressive modelling to exploit the predictability of information in one brain area by past activity in another (*39*). We fit GPDC models to the 600 ms that began 300 ms after the state cue onset, which corresponded to the period of rising theta coherence. Models were fit in both directions (HPC→OFC, OFC→HPC) and the difference was taken to obtain the net signal directionality. Significance was assessed by measuring whether the bootstrapped confidence intervals of the net signal directionality overlapped with zero.

### Phase modulation of single neurons

Phase-modulation of HPC and OFC single neurons was assessed by examining the distribution of a given neuron’s spiking with respect to the phase angle of the simultaneously recorded LFP. Instantaneous LFP phase angles were determined by bandpass filtering the LFPs to a frequency of interest (theta (4–8 Hz), alpha (9-12 Hz), beta (13–30 Hz), and gamma (30–60 Hz)) and then extracting phase information via the Hilbert transform. We used Rayleigh’s test of circular uniformity to assess the extent to which spikes clustered at a specific phase angle as compared to the same number of spikes being uniformly distributed across all phase angles.

Theta peaks during the state epoch were identified by first bandpassing the HPC and OFC LFP channels in the theta band (4-8 Hz). Next, we extracted the amplitude of the bandpassed LFP via the Hilbert transform. We then z-scored theta amplitudes on each channel and aligned them to the onset of the state cue within each individual trial. We defined each trial’s theta peak as the first positive peak of the single-trial theta-amplitudes after state-cue onset that was greater than the mean (z > 0). Only recording sites that had at least one well-isolated neuron were included in the analysis.

### Statistics

All statistical tests are described in the main text or the corresponding figure legends. Error bars and shading indicate 95% confidence intervals unless otherwise specified. We used a bootstrapping procedure to estimate confidence intervals. This procedure involved randomly subsampling 80% of the data and then computing the relevant statistic. This process was repeated 1000 times, each time randomly subsampling the data. The 5^th^ and 95^th^ percentiles of the resulting distributions were then used as confidence intervals. All terms in regression models were normalized and had maximum variance inflation factors of 1.7. All comparisons were two-sided.

## Data and code availability

The dataset and code supporting the current work are available from the corresponding author on request.

## Notes

### Competing Interest Statement

The authors have declared no competing interest.

